# Photoacoustic imaging reveals transient decrease of oxygenation in murine blood due to monoclonal IgG4 antibody

**DOI:** 10.1101/2022.09.29.509334

**Authors:** Anjul Khadria, Chad D. Paavola, Konstantin Maslov, Patricia L. Brown-Augsburger, Patrick F. Grealish, Emmanuel Lozano, Ross L. Blankenship, Rui Cao, Junhui Shi, John M. Beals, Sunday S Oladipupo, Lihong V. Wang

## Abstract

Over 100 monoclonal antibodies have been approved by the FDA for clinical use; however, a paucity of knowledge exists regarding the injection site behavior of these formulated therapeutics, i.e., the effect of antibody and formulation on the tissue around the injection site and vice versa. In this report, we injected a near-infrared dye-labeled IgG4 isotope control antibody into the subcutaneous space in mouse ears to analyze the injection site dynamics, including quantifying molecular movement. Surprisingly, we discovered that the antibody reduces the local blood oxygen saturation levels in mice after prolonged anesthesia without affecting the total hemoglobin content and oxygen extraction fraction. The local oxygen saturation results open a new pathway to study the functional effects of monoclonal antibodies.

## Main

Monoclonal immunoglobulin G (IgG) antibody-based drugs have provided lifesaving options for people with various diseases, including COVID-19 [1,2]. Over the last four decades, over 100 monoclonal antibodies have received FDA approval to treat several disorders, with a preponderance approved in the previous decade [3,4]. Despite being extensively utilized as therapeutics, limited knowledge exists regarding their behavior at the injection site, which may hinder development and optimal molecule selection [5,6]. Following subcutaneous injections, monoclonal antibodies are slowly absorbed via the lymphatic system due to their large hydrodynamic size; however, the formulation excipients can diffuse away from the injection site rapidly via absorption across the microvasculature [7,8]. The slow absorption of the antibodies from the injection site via the lymphatic system extends the residence time at the injection site; thus, the behavior of these molecules is ultimately affected by the local conditions of the injection site, i.e., the pH, temperature, architecture of extracellular matrix, viscosity of the interstitial fluid, and muscle movement [9–12]. Apart from being affected by the injection site conditions, the antibodies themselves can influence changes in tissue characteristics, e.g., temporary hypersensitivity reactions [13,14]. The study of changes in local hemodynamic features such as oxygen saturation (oxygenation), oxygen extraction fraction (OEF), and change in total hemoglobin could provide insights into unknown effects of antibodies at the injection site. Apart from increasing use as drug therapeutics, monoclonal antibodies conjugated with contrast agents have found increased use for visualization of different structures in the body through different imaging modalities [15–17]. Thus, the study of the effects of antibodies on the injection site characteristics may further help in designing safer and more effective therapies and reagents for medical use.

Here, we use optical-resolution photoacoustic microscopy (OR-PAM) to study the subcutaneous injection site dynamics of near-infrared (NIR) dye-labeled IgG4 isotype control antibody, as well as its effects on the local hemodynamics at the injection site of mice ears. We selected IgG4 isotype control antibody with amino acid substitutions to minimize effector function as a testbed for this work due to its ready availability, favorable biophysical properties, and lack of target mediated interaction at the site of injection. While the variable domains of this antibody do not have any specific paratope, the constant domain of the antibody is still able to engage in binding to Fc receptors during pinocytosis [18,19].

## Results and Discussion

### Injection site dynamics of antibody

We labeled IgG4 antibody with the sulfo-cy7.5 dye through a previously described method [17]. Before analyzing the absorption behavior of the dye-labeled IgG4 antibody by photoacoustic imaging, we first investigated the suitability of the mouse ear model for this study, as well as the effects of labeling on the pharmacokinetics of the antibody. Dye-labeled and unlabeled IgG4 isotype control antibody solutions were injected in mice by the intravenous route and by subcutaneous injections in the torso and ear. Pharmacokinetic samples were collected over 504 hours (21 days) following administration (Figures 1a and S1a). We observed similar pharmacokinetic profiles for both subcutaneous locations, thus confirming that the mouse ear model is suitable for our study. We also demonstrated that cy7.5 dye labeling did not appreciably alter the pharmacokinetic properties of the antibody by comparing the pharmacokinetic profiles of the dye-labeled and unlabeled antibody solutions in the mouse ear (Figure 1b).

**Figure 1:**
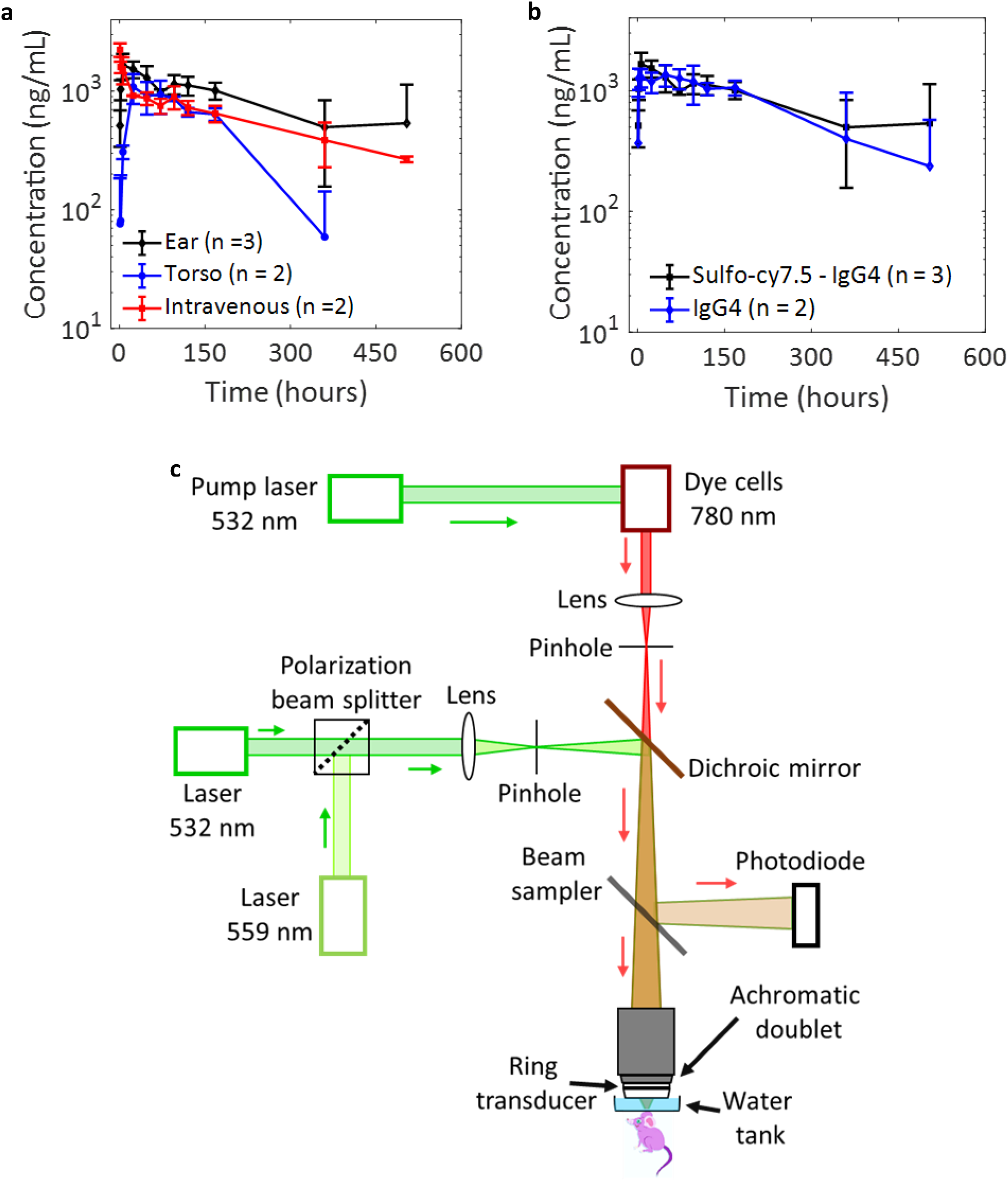
(a) Comparison of bioavailability of the dye-labeled antibody through different routes of injections in mice. (b) Comparison of dye-labeled and unlabeled antibody dosed subcutaneously in mice ears shows that the dye labeling does not alter the pharmacokinetics of the antibody. (c) Schematic of the triple-wavelength equipped OR-PAM. All data represent mean ± standard deviation.

We designed and employed a triple-wavelength equipped OR-PAM (Figure 1c) to study the near-infrared light absorbing sulfo-cy7.5 dye-labeled IgG4 isotype control antibody in the mouse ear. Our initial efforts to quantify the injection site absorption kinetics of the dye-labeled IgG4 antibody through photoacoustic imaging using the previously reported method were unsuccessful (SI, Figures S1b and S1c) [20]. The results suggested less than 25% of the dye-labeled antibody disappeared from the injection site after the first 3 hours; however, at 6 hours and 24 hours’ time points, the total photoacoustic signal was higher than the total signal just after injection (i.e., at 3 minutes), which could not be the true reflection of the injection site kinetics of the antibody. The increase in the total photoacoustic signal was hypothesized to be the result of changes in the environment of the dye-labeled antibody, which led to a dynamic change in its molar absorption coefficient. In such a scenario, a change in photoacoustic signals cannot be directly considered as the change in the concentration of dye-labeled antibody [21]. The antibody solution was prepared in phosphate-buffered saline (PBS), which can be readily absorbed across the microvasculature, while the hydrodynamically large antibody can only be absorbed via the lymphatic vessels at a concomitantly slower rate due to diffusion and drainage of the lymphatic fluids [7,22]. Since the salts in the PBS vehicle are absorbed at a different rate than the antibody, the changing solution conditions in the injection site result in dynamic changes to the molar absorption coefficient of the dye-labeled antibody, making quantification studies unreliable.

Pharmacokinetic quantification of dye-labeled antibodies prepared in buffer solutions using fluorescence microscopy has been reported several times [23–25]. However, due to the dynamic change of molar absorption coefficient due to different absorption mechanisms of the carrier and the antibody, most of the studies have observed an increase in fluorescence, thus impacting the pharmacokinetic analysis. As a negative control, we quantified the absorption kinetics of sulfo-cy7.5 dye alone, dissolved in the PBS buffer, and observed that most of the dye (∼ 90%) disappears from the injection site within the first 3 hours of injection (SI, Figure S1d).

We studied the antibody movement at the injection site during imaging by estimating the change in the lateral 2D area occupied by the dye-labeled antibody. In the first 60 minutes, the area occupied by the antibody increased by 10 – 15% (Figure 2c). This is in stark contrast to a smaller size dye-labeled insulin lispro (∼ 6.8 kDa) in Humalog formulation, whose area was reported to increase by 50 – 60% in the first 60 minutes using the same photoacoustic imaging technique [20]. After 24 hours, the area occupied by the antibody is roughly four times the initial area following injection (Figure 2d). At 3 hours post-injection, when a majority of the PBS is absorbed by the blood vessels, the antibody is primarily dissolved in the interstitial fluid (SI, Figure S2d), and hence, the flow of interstitial fluid may significantly influence the movement of the antibody apart from other factors such as diffusion and convection [5,26,27].

**Figure 2:**
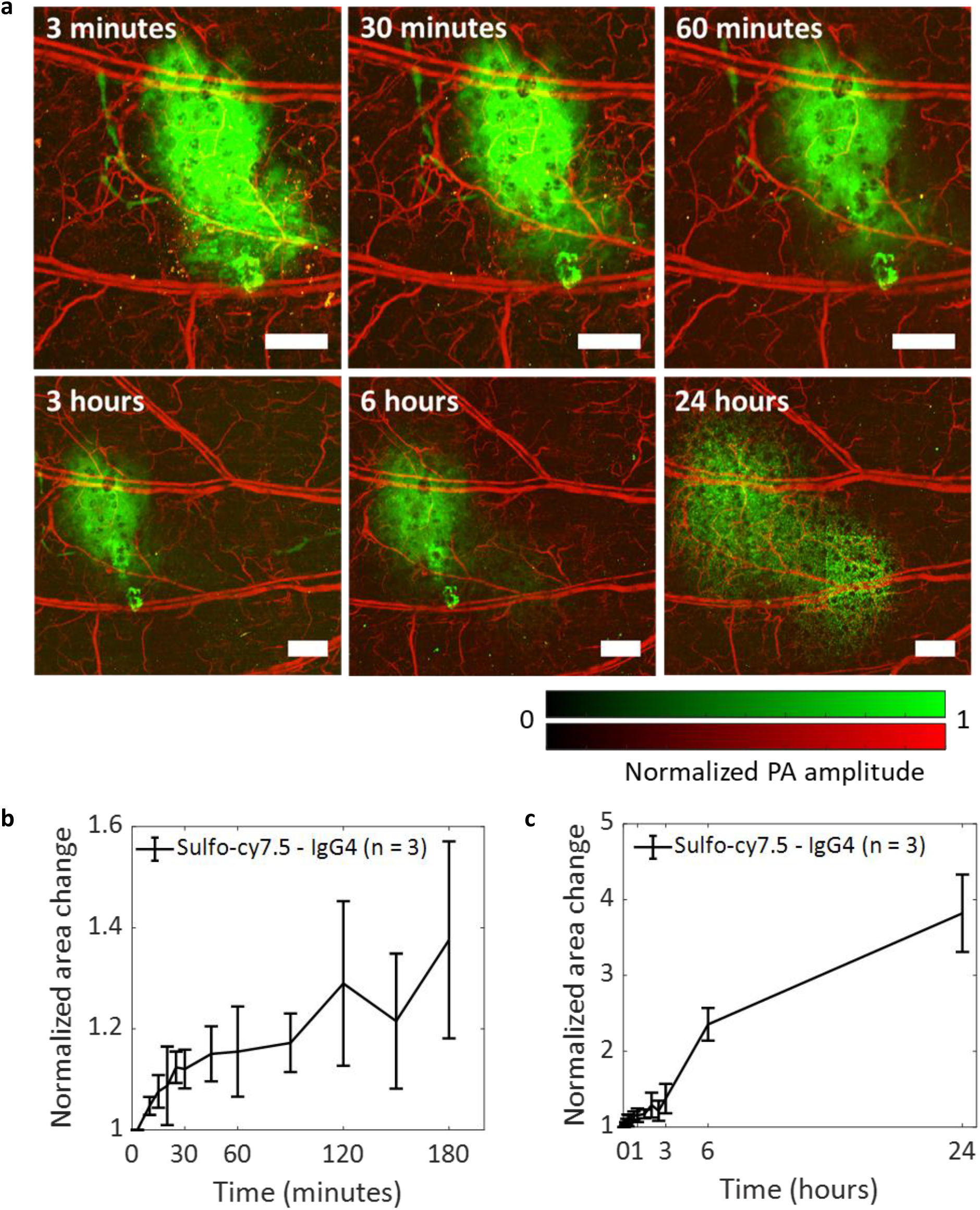
Diffusion of the dye-labeled IgG4 antibody. (a) OR-PAM shows diffusion of dye-labeled IgG4 antibody in the mouse ear. The images at 3 hours, 6 hours, and 24 hours are of a larger field of view. (b) The area of the sulfo-cy7.5 dye labeled IgG4 antibody increased by about 40% after the first 3 hours after injection. (c) The antibody area increased by about 300% after 24 hours of injection. The mouse was awake and moving between 3 hours and 6 hours, and 6 hours and 24 hours’ time points. All data represent mean ± standard deviation (n = 3). PA, photoacoustic. Scale bars, 500 μm.

### Measurement of oxygen saturation (sO_2_) in local blood

We monitored the effects of the dye-labeled IgG4 isotype control antibody on the local blood sO_2_ through photoacoustic imaging using the 532 nm and 559 nm light [28]. After subcutaneous injection of the dye-labeled antibody, we initially saw an increase in local blood sO_2_ in veins due to the needle insertion. However, approximately 2 hours post-injection, the value of sO_2_ in both the veins and arteries dropped (Figure 3a). After switching off the isoflurane supply, and keeping the mouse awake, the sO_2_ returned to the normal physiological levels without observation of any local or systemic adverse effects on the mouse. Notably, we did not observe any sO_2_ decrease upon injecting sulfo-cy7.5 in PBS (Figure 3b). However, like the antibody, the injection of sulfo-cy7.5 in PBS instantaneously led to the increase of venous sO_2_ at the point of injection to over 95%, which then gradually decreased over time to return to normal physiological levels [29]. The insertion of an empty sterile needle produced a similar result; thus, suggesting that the sO_2_ increase is not due to the chemical effects of the formulations or buffers, but likely due to the needle. It is not fully understood what mechanism may result in the sudden and significant rise of local blood sO_2_ levels, which takes a long time to return to normal physiological levels in the case of sulfo-cy7.5 or only a sterile needle (Figure 3c). The hypothesis is the needle might be causing damage to the tissue leading to transient local tissue hypoxia [30,31] and hence, the high sO_2_ blood from arteries is directly passed into the veins. However, in the antibody studies, although the value of local blood sO_2_ decreased after 2 hours (Figure 3d), the total local hemoglobin amount remained similar (Figure 3e) for the duration of the experiment; thus, indicating that there is no change in the local blood content. We did not observe any significant change in the local OEF 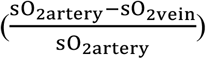 indicating that the oxygen consumption of the tissue is unlikely to play a role in the decrease of local blood sO_2_ (SI, Figure S2a). To exclude the possibility that anesthesia used in the imaging experiments is responsible for this, we imaged a mouse ear for five hours without performing any injection and observed no changes in the local blood sO_2_ (SI, Figure S2b). The observed local blood sO_2_ decrease for unlabeled IgG4 isotype control antibody confirmed that the sulfo-cy7.5 dye labeling is unlikely to play any role (SI, Figure S2c). To verify if the decrease of local blood sO_2_ is due to the combination of the monoclonal IgG4 antibody and anesthesia, we performed an experiment wherein, the isoflurane was switched on and off at regular intervals (Figure 4). Upon switching off the isoflurane for 10 minutes after 3 hours of anesthesia, we observed an increase in the sO_2_ levels across the whole field of view, which returned to normal physiological levels within the next 10 – 15 minutes. Upon re-anesthetizing with isoflurane, the sO_2_ decreased again after about 2 – 2.5 hours and surged again upon switching off the isoflurane. The mouse was then fully awakened and left to freely roam in the cage (with food and water) for around 2 – 2.5 hours. The muscle movement in the ears during this period was hypothesized to facilitate lymphatic absorption of the antibody. Upon re-anesthetizing and re-imaging, the same mouse, the sO_2_ decreased to a lesser extent, over a longer period, requiring up to 5 hours as opposed to 2 – 2.5 hours, and rose again after switching off the isoflurane. We observed a similar pattern upon keeping the mouse awake in its cage for 10 hours. While the changes in sO_2_ were striking, we did not observe any acute adverse effects or discomfort in any of the mice that were injected with the labeled antibody.

**Figure 3:**
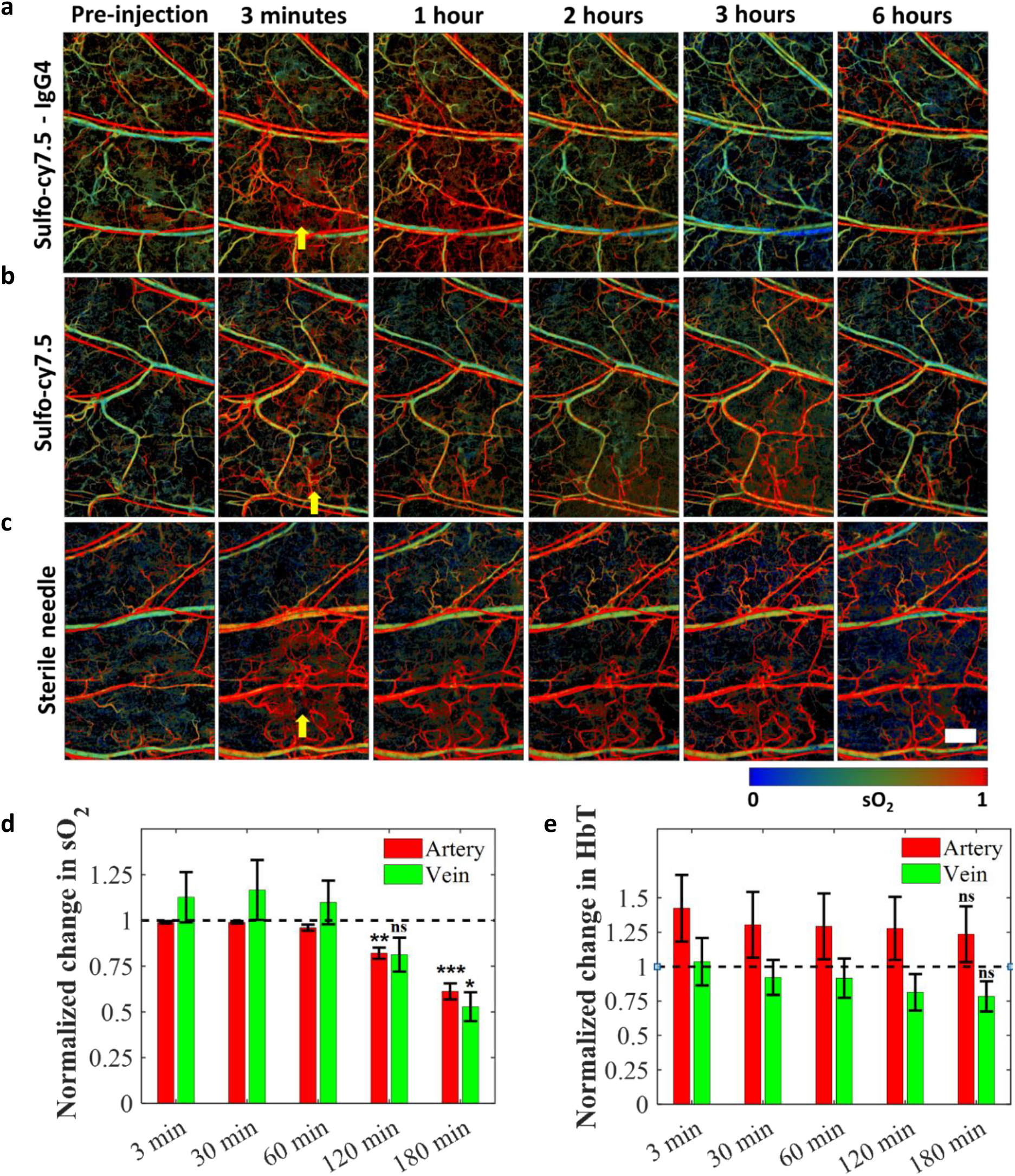
Functional changes in the mouse blood caused by the antibody. (a) Blood sO_2_ starts decreasing after 2 hours of injection of the dye-labeled IgG4 antibody. (b) No significant change in blood sO_2_ after injection of sulfo-cy7.5 dye in PBS. (c) Insertion of sterile needle does not cause any decrease in the blood sO_2_. (d) Statistically significant changes in sO_2_ of major arteries and veins are observed with respect to the pre-injection levels. (e) No significant change in the total hemoglobin (HbT) was observed throughout imaging. The horizontal dashed lines refer the pre-injection values. Number of mice, n = 5; data represent mean ± standard error of the mean. All p values at the mentioned time points were calculated using paired t-test with respect to the pre-injection time points; p > 0.05, ns; p < 0.05, *; p < 0.01, **; p < 0.001, ***. Yellow arrows represent points of injections. The mice were kept awake between 3 hours and 6 hours time points. Scale bar, 500 μm.

**Figure 4:**
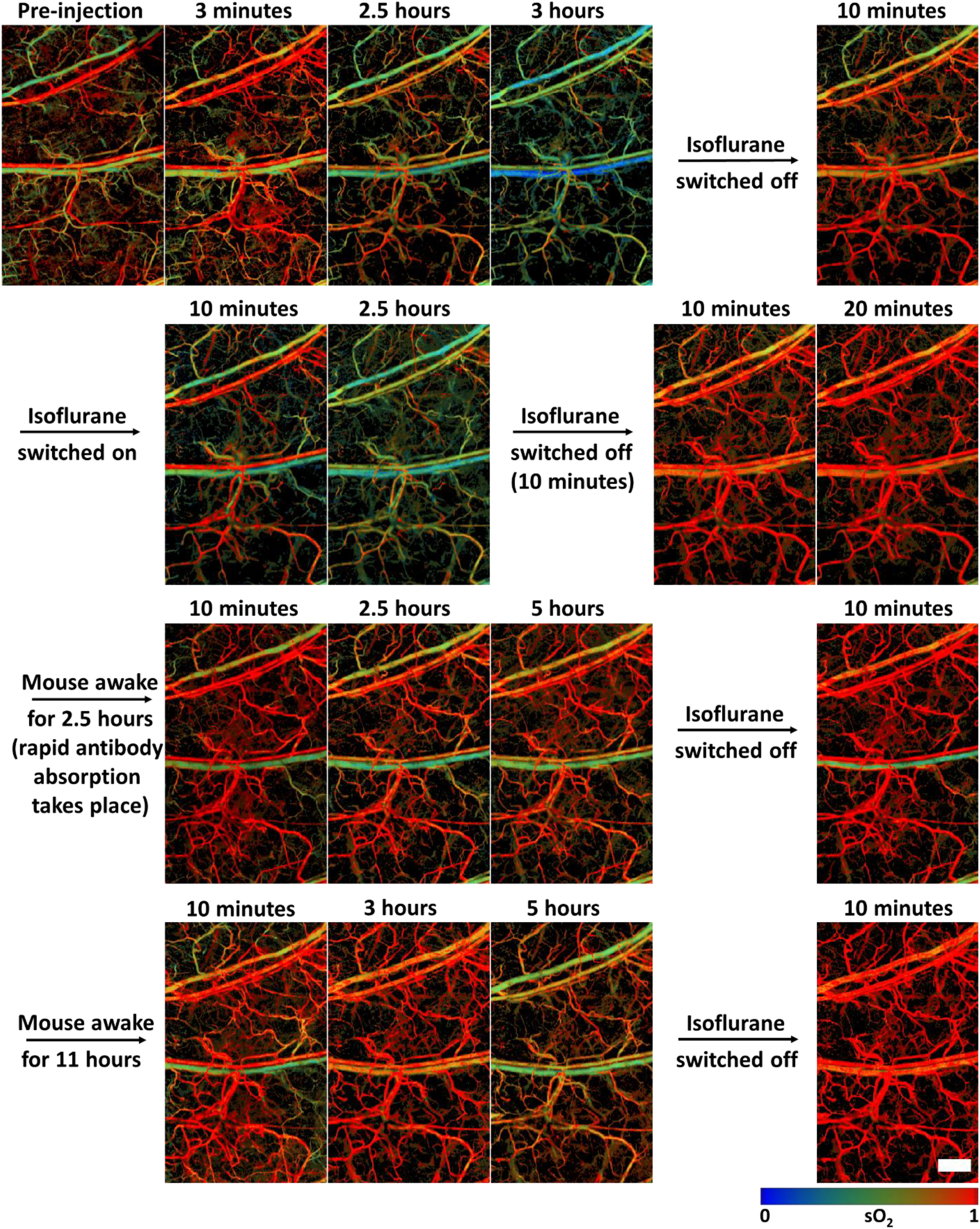
Dependence of blood sO_2_ on isoflurane anesthesia upon injection of dye-labeled IgG4 antibody in a mouse. The time points are reset to zero at every isoflurane on/off activity. Scale bar, 500 μm.

### Conclusions

Through pharmacokinetic experiments of dye-labeled and unlabeled antibody, we show that the extent of dye labeling in these experiments did not affect the absorption characteristics of the antibody, and that subcutaneous administration in the mouse ear was suitable to study the subcutaneous behavior of the monoclonal IgG4 antibody. Our experiments prove that light-based optical absorption techniques cannot quantify injection site absorption of large-sized molecules such as antibodies unless the dynamic change in the molar absorption coefficient is deduced. We also quantified antibody movement at the injection site caused by interstitial fluid flow. Through a series of experiments, we show that monoclonal human IgG4 antibody locally reduces oxygen saturation in mouse blood after prolonged anesthesia, and that oxygenation is recovered within a few minutes after switching off the isoflurane. The underlying mechanisms that lead to oxygenation decrease in anesthetized mice are not well-understood at this point and further experiments are required, which are beyond the scope of this work. Given the widespread preclinical and clinical use of antibodies at an exponentially increasing rate, these results open new pathways and call for further investigations of their injection site characteristics and effects.

## Methods

The sulfo-cy7.5 dye-labeled IgG4 antibody was prepared and characterized as previously reported [17].

### Optical resolution photoacoustic microscopy (OR-PAM) system design

We used a ring-shaped transducer (central frequency = 42 MHz, f-number = 1.67, Capistrano Labs) in the OR-PAM setup, which is equipped with three lasers of 532 nm, 559 nm, and 780 nm optical wavelengths. Light pulses of all the three wavelengths were used to shine at the same point successively with microseconds delay. The light beams from a 559 nm Nd:YAG laser (BX2II, Edgewave) and 532 nm light (SPOT-10-200-532, Elforlight) laser were combined using a polarization beam splitter (PBS121, Thorlabs) and focused on a 254 μm diameter orifice (3928T991, McMaster-Carr) for spatial filtering. The 780 nm light was emitted from a dye (Styryl 11 dye in 200 proof ethanol, 07980, Exciton) laser (Credo, Sirah) pumped by a 532 nm Nd:YAG laser (IS80-2-L, Edgewave GmbH). The 780 nm beam was spatially filtered by focusing it on a second pinhole (3928T991, McMaster-Carr). The 532 nm and 559 nm light beams were combined with the 780 nm light beam through a dichroic mirror (M254C45, Thorlabs). The combined light beam was focused on the sample through an achromatic doublet (AC080-020-A, Thorlabs) after collecting some amount of light by a photodiode (PDA36A, Thorlabs) through a beam sampler to correct for laser fluctuations. The mouse was scanned using a two-dimensional scanner built with stepper motors (PLS-85, Physik Instrumente) that were controlled by a customized LabVIEW program with an FPGA (PCIe-7841, National Instruments). All the data was acquired through a digitizer (ATS 9350, AlazarTech) at a frequency of 500 MS/s.

### Animal experiments

#### Imaging experiments

We performed all the imaging experiments on animals using protocols approved by IACUC at Caltech. We used Hsd:Athymic Nude-Fox1^nu^ mice aged 6 to 12 weeks (Envigo) in all the photoacoustic experiments while maintaining their body temperatures at 37 °C during imaging. The mice were acclimated for at least 4 days before imaging experiments. All the mice were imaged under isoflurane anesthesia (1.25 – 1.50 % isoflurane in the air at a flow rate of 1L/min).

#### Pharmacokinetic studies of human IgG4 and sulfo-cy7.5 labeled human IgG4 antibodies

Mouse pharmacokinetic study protocols were approved by the IACUC at Eli Lilly and Company. Female CD-1 mice (6 to 12 weeks old, approximately 20 – 35 g in weight) were obtained from Envigo (Indianapolis, IN). Animals were acclimated for at least 4 days before test article administration. Unlabeled IgG4 and the sulfo-cy7.5 dye-labeled IgG4 antibody samples were formulated at 0.02 mg/mL for intravenous (IV) and torso subcutaneous (SC) dosing or 20 mg/mL for SC ear dosing in phosphate buffered saline (PBS), pH 7.2. Mice were anesthetized using isoflurane by inhalation of approximately 2% in the air. A volume of 200 mL of the 0.02 mg/mL antibody was delivered by the IV (tail vein) or SC (torso) route (0.004 mg/mouse). For SC administration in the ear, a volume of 0.2 μL was administered using a 2.5 microliter glass syringe (600 Series, Hamilton) with removable needle assembly (7632-01; Hamilton) affixed with a 34-gauge, 0.5-inch needle (207434; Hamilton) (0.004 mg/mouse). Sample collection was performed by making sequential tail nicks at 1, 2, 6, 24, 48, 72, 96, 120, 168, 336, and 504 hours after test article administration. 10 μL of whole blood was collected with a microcapillary (Drummond Scientific), and the sample was then immediately dispensed into 90 μL Reagent E buffer (Gyros, Uppsala Sweden). The resulting 100 μL sample was centrifuged (2000-RCF, 10 minutes), after which the supernatant was transferred to a labeled polypropylene cluster tube and stored at −70 °C before analyses. Human IgG concentrations were determined in the whole blood samples performed on the Gyros xPand instrument using the Gyros Generic hIgG pharmacokinetic kit (P0020499). The standard curve range was from 2 to 2000 ng/mL for IgG or 5 to 5000 ng/mL for sulfo-cy7.5 dye-labeled IgG4.

Standard curve regression was performed on Gyros Evaluator regression software to interpolate unknown sample concentrations using a five-parameter logistical fit model of the fluorescence responses at the 1%-photomultiplier tube (PMT) setting (Gyrolab User Guide, 2018; section D, Data Analysis, Chapter 4, p. D-33)

The concentrations determined based on the 10% mouse blood/90% Rexxip A buffer samples were transformed to plasma concentrations by multiplying by a factor of 17.36. The correction factor accounts for mouse hematocrit and dilution effects during whole-blood sample processing [32].

Plasma pharmacokinetic parameters were determined using Phoenix WinNonLin version 8.1.0.3530. Values below the quantification limits were ignored in the pharmacokinetic parameter calculations.

### Imaging protocol

We adapted an imaging protocol as previously reported.

### Calculating the area occupied by the dye-labeled IgG antibody bolus to study its movement

Maximum amplitude projection (MAP) images of the sulfo-cy7.5 dye-labeled IgG4 antibody (0.1 μL, 20 mg/mL, n = 3) were acquired from the raw photoacoustic data. The MAP images were thresholded (after passing through a median filter of size 4 × 4 pixels) by the summation of the mean and three times the standard deviation of the background amplitude to segregate photoacoustic signals (generated by the dye-labeled antibody) from the noise in the region of interest. The total number of pixels within the thresholded region of interest was multiplied by the size of a single pixel (2.5 μm x 5.0 μm) to calculate the area occupied by the antibody bolus. The area occupied at each time point was divided by the area at 3 minutes (just after injection) to calculate the normalized area.

## Data availability

The data that support the conclusions are mentioned in the main draft or the supplementary information.

## Author contributions

L.V.W., S.O., J.M.B., C.D.P., and A.K. conceived the project and the ideas. C.D.P. and A.K. designed the chemistry and parameters for dye labeling. P.G. labeled the antibody with the dye and characterized them. P.G. and A.K. prepared the antibody and dye buffer solutions. A.K. and K.M. designed and built the scanning photoacoustic microscope. A.K. designed and performed all the photoacoustic experiments and analyzed all the photoacoustic data. R.C. and J.S. wrote the LabVIEW software for photoacoustic data acquisition. P.B.A., E.L., and R.L.B. designed, performed, and analyzed the pharmacokinetic experiments. L.V.W., S.O., and J.M.B. supervised the project. A.K. wrote the manuscript. C.D.P., P.B.A., J.M.B, S.O., and LV.W. contributed to writing the manuscript.

