## Supplementary info for "Photoacoustic imaging reveals transient decrease of oxygenation in murine blood due to monoclonal IgG4 antibody"

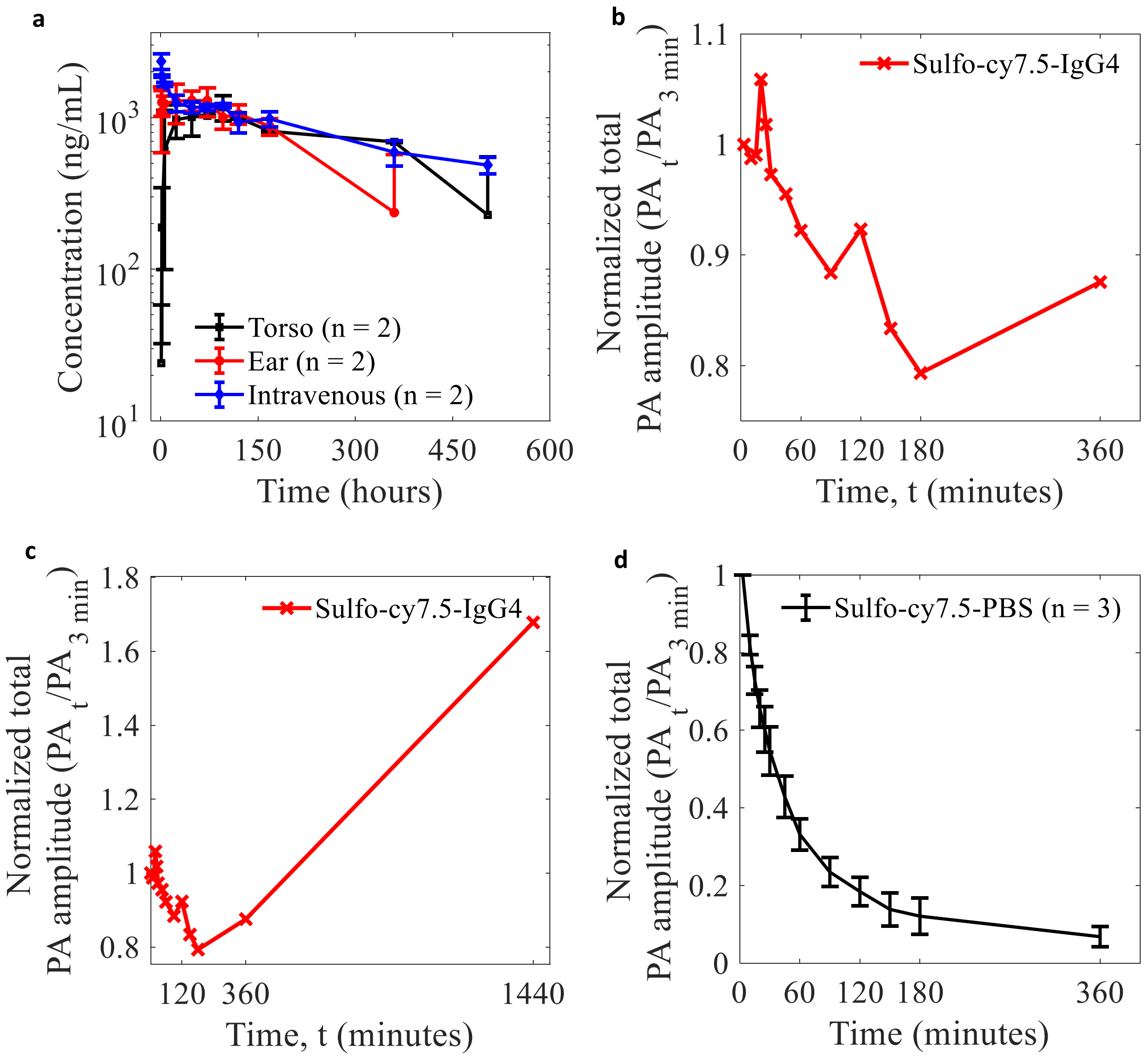


**Figure S1: Quntification.** (a) Pharmacokinetics data of unlabeled IgG4 isotype control antibody in mice following IV and subcutaneous administration in the torso or ear; data represent mean ± standard deviation. (b, c) Photoacoustic quantification of absorption of dye-labeled IgG4 antibody for 24 hours and 6 hours. (d) Photoacoustic quantification of sulfo-cy7.5 dye dissolved in PBS buffer; data represent mean ± standard error of the mean.

**Table S1: Mean pharmacokinetic parameters of unlabeled IgG4 antibody**

| | $t_{1/2}$ (h) | $T_{max}$ (h) | $C_{max}$ (ng/mL) | $AUC_{0-168}$ (h*µg/mL) | $AUC_{all}$ (h*µg/mL) | $AUC_{INF\_obs}$ (h*µg/mL) | Cl_obs (mL/h/kg) |
|---|---|---|---|---|---|---|---|
| **IV-tail** | 304 | 1 | 2351 | 195 | 414 | 629 | 0.26 |
| **SC-torso** | 520 | 96 | 1179 | 160 | 408 | 867 | 0.20 |
| **SC-ear** | 511 | 48 | 1355 | 197 | 330 | 944 | 0.19 |

**Table S2: Mean pharmacokinetic parameters of sulfo-cy7.5 dye-labeled IgG4 antibody**

| | $t_{1/2}$ (h) | $T_{max}$ (h) | $C_{max}$ (ng/mL) | $AUC_{0-168}$ (h*µg/mL) | $AUC_{all}$ (h*µg/mL) | $AUC_{INF\_obs}$ (h*µg/mL) | Cl_obs (mL/h/kg) |
|---|---|---|---|---|---|---|---|
| **IV-tail** | 238 | 1 | 2233 | 145 | 288 | 379 | 0.40 |
| **SC-torso** | 117 | 36 | 1206 | 130 | 158 | 235 | 0.70 |
| **SC-ear** | 444 | 12 | 1808 | 199 | 405 | 1005 | 0.30 |

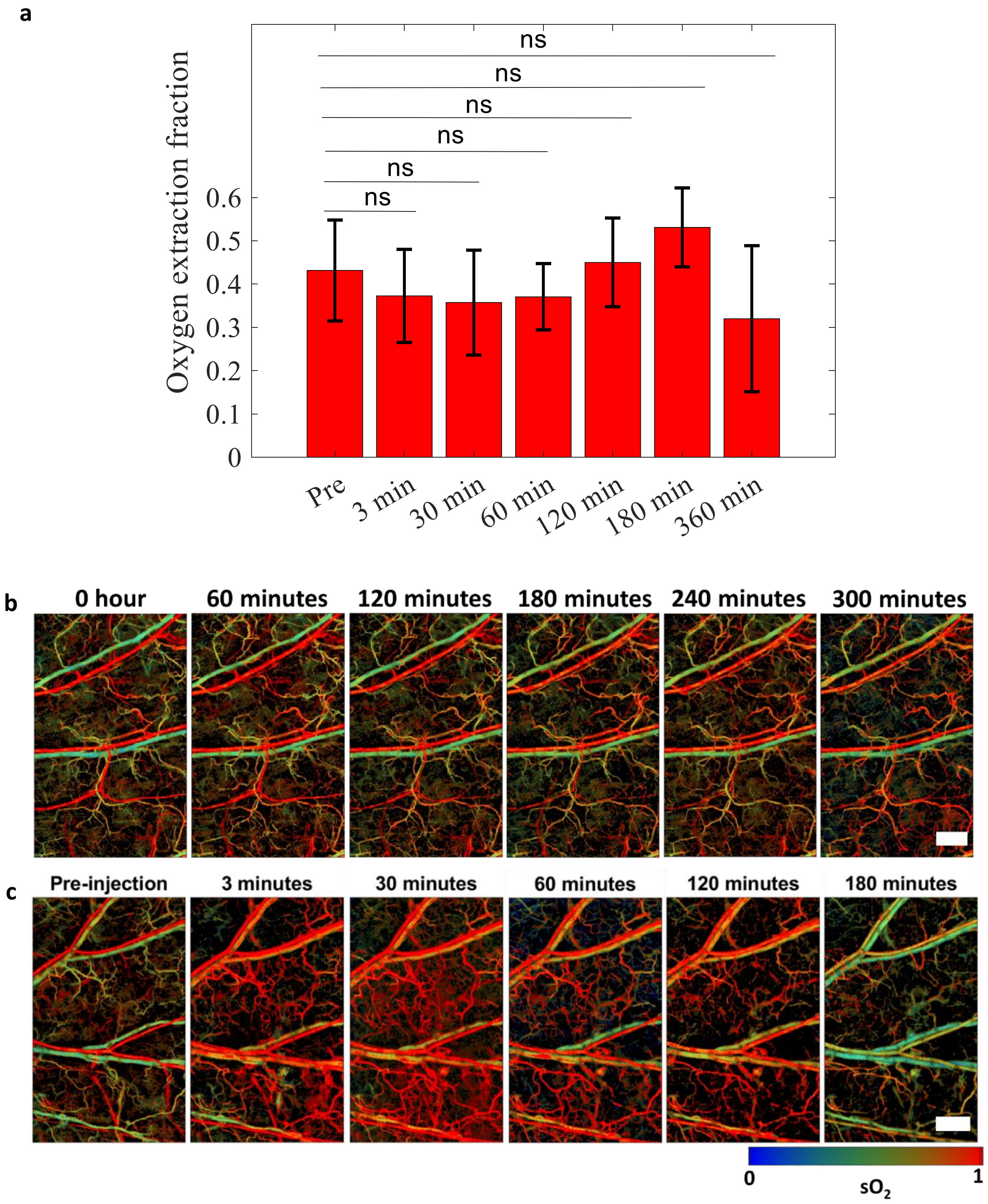


**Figure S2: Functional changes caused by the antibody.** (a) No significant change cuased by the dye-labeled antibody in the oxygen extraction factor with respect to the preinjection value. Number of mice, n = 5; data represent mean ± standard deviation. All p values at the mentioned time points were calculated using paired t-test with respect to the pre-injection time point; $p > 0.05$, ns. (b) No change of blood $sO_2$ in mouse ear after being kept under anesthesia for 5 hours continuously. (c) Blood $sO_2$ in the mouse ear decreases after injection of unlabeled IgG4 antibody. Scale bars, 500 μm.
